# Molecular and functional heterogeneity of cerebellar granule cell terminals expands temporal coding in molecular layer interneurons

**DOI:** 10.1101/338152

**Authors:** Kevin Dorgans, Valérie Demais, Yannick Bailly, Bernard Poulain, Philippe Isope, Frédéric Doussau

**Author notes:** Corresponding author: Frédéric Doussau, INCI, CNRS UPR 3212, 5, rue Blaise Pascal, 67084 Strasbourg, France.

## Abstract

In the cerebellum, molecular layer interneurons (MLIs) play an essential role in motor behavior by exerting precise temporal control of Purkinje cells, the sole output of the cerebellar cortex. The recruitment of MLIs is tightly controlled by the release of glutamate from granule cells (GCs) during high-frequency activities. Here we study how single MLIs are recruited by their distinct unitary GC inputs during burst of GC stimulations. Stimulation of individual GC-MLI synapses revealed four classes of connections segregated by their profile of short-term plasticity. Each class of connection differentially drives MLI recruitment. Molecular and ultrastructural analyses revealed that GC-MLI synaptic diversity is underlain by heterogeneous expression of synapsin II at individual GC terminals. In synapsin II knock-out mice, the number of classes is reduced to profiles associated with slow MLI recruitment. Our study reveals that molecular diversity across GC terminals enables diversity in temporal coding by MLIs and thereby influences the processing of sensory information by cerebellar networks.

## Introduction

Inhibitory interneurons mediating feed-back or feed-forward inhibition (FFI) provide brain microcircuits with an exquisite temporal control over the firing frequency of projecting neurons ^1–4^. In the cerebellar cortex, the FFI microcircuit is activated by granule cells (GCs) that target two types of molecular layer interneurons (MLIs), stellate cells (SCs) and basket cells (BCs). SCs and BCs finally control Purkinje cells, the sole projecting neurons of the cerebellar cortex, through a powerful somatic or dendritic inhibition ^5^ In combination with the direct excitatory pathway provided by GC-PC connections, the FFI encodes sensorimotor information through acceleration or deceleration of PC simple spike activity ^6–8^.

Sensorimotor information is conveyed to the cerebellar cortex by mossy fibers (MFs) as short high-frequency bursts of action potentials ^9–13^. During high-frequency stimulations, cerebellar synapses exhibit several forms of short-term synaptic plasticity (STP) including facilitation and depression of synaptic responses in the millisecond range ^14–21^. STP play multiple roles in information processing ^22,23^; in the cerebellar cortex, differences in the profile of STP across synapses involved in the direct excitatory pathway or in the FFI microcircuit control the inhibitory/excitatory balance and shape Purkinje cell discharge ^24^. GCs, which are the most numerous neurons in the brain, are the key actors of the inhibitory/excitatory balance in the cerebellar cortex, but STP heterogeneity across GC boutons is poorly documented. A seminal study has shown that the behavior of glutamate release during high-frequency activities at GC boutons is determined by the target cell (that is PC, SC or BC, Bao et al., 2010). Following compound stimulations of clusters of GCs or beams of parallel fibers (PFs), it was shown that GC-BC synapses depress during high-frequency stimulation while GC-SC synapses facilitate ^19^. By controlling the spatiotemporal excitability of PC ^19^, and potentially by shaping the inhibitory/excitatory balance ^24^, target cell–dependency of STP at the input stage of the FFI pathway must have important functional consequences for cerebellar output. However, how synaptic heterogeneity relates to target-cell specificity remains unknown. Besides, target cell–dependency of STP at GC-MLI synapses is challenged by different experimental findings. First, many MLIs cannot be classified solely by their axon profile (*e.g*. basket versus dendritic synapses) or their position in the molecular layer ^25,26^ It was suggested that MLIs represent a single population of interneurons ^5,25,27,28^ Second, release properties and STP profiles of GC synaptic inputs to MLIs can be modified by presynaptic long-term plasticity and by local retrograde signaling independently of the target cell ^29,30^. Given the abundance of GCs and the importance of STP at GC-MLI synapses for cerebellar computation, we set out to study the diversity of STP at unitary GC-MLI synapses.

We show that MLIs, regardless of their identity, receive at least four functionally distinct types of synapses from GCs. Each class of connection differentially drives the timing of MLI recruitment characterized by the first-spike latency. Differences in STP across GC-MLI connections are determined by distinct ultrastructural properties and molecular contents of GC boutons. We reveal that synapsin II (Syn II), a presynaptic protein involved in STP ^31,32^, is heterogeneously expressed at GC-MLI synapses. Functional studies using wild-type (WT) and Syn II KO mice demonstrate that Syn II determines the profile of STP and the first-spike latency in MLI. Our observation that single MLIs receive GC inputs with distinct molecular and functional properties suggests that the temporal coding of GC activity by MLIs and thereby the inhibitory control over the cerebellar output via PCs depends on GC subtypes recruited by a given sensorimotor input.

## RESULTS

### Functional heterogeneity at unitary GC-MLI synapses during high-frequency stimulations

In order to study how information from single GC inputs is encoded by MLIs, we measured STP at unitary GC-MLI synapses in acute parasagittal slices by minimal electrical stimulation of PFs (10 pulses at 100Hz), in the direct vicinity of the dendritic tree of a recorded MLI ^18,33^ MLIs localized in the vermis (lobules IV-VI) were recorded in whole-cell voltage-clamp configuration and loaded with Alexa-594 (n = 49) to visualize their dendritic tree and their morphology. Using two-photon microscopy, the stimulation pipette was visually positioned above an isolated dendrite. MLI subtypes were identified in half of the recorded cells based on the presence of a basket in the Purkinje cell layer (n = 13 for BC, n= 12 for SC and n = 24 for non-identified MLI). In 28 MLIs, we recorded synaptic responses from at least 2 GCs. STP profiles were highly heterogeneous across unitary GC-MLI contacts (n = 115) including unitary connections contacting the same MLI (***Figure 1***). To classify the STP profiles, we used principal component analysis (PCA) on the averaged and normalized synaptic charges in trains of EPSCs followed by a *k*-means clustering analysis (***Supplementary figures 2,3*).** We identified 4 clusters that characterize STP at GC-MLI synapses (***Figure 2A-B*).** The profiles differ by: (i) the quantity of neurotransmitter released at the first stimuli (***Figure 2C***) (ii) the STP profiles during the first four EPSCs, and (iii) the ability to sustain glutamate release after the fourth stimuli.

**Figure 1.**
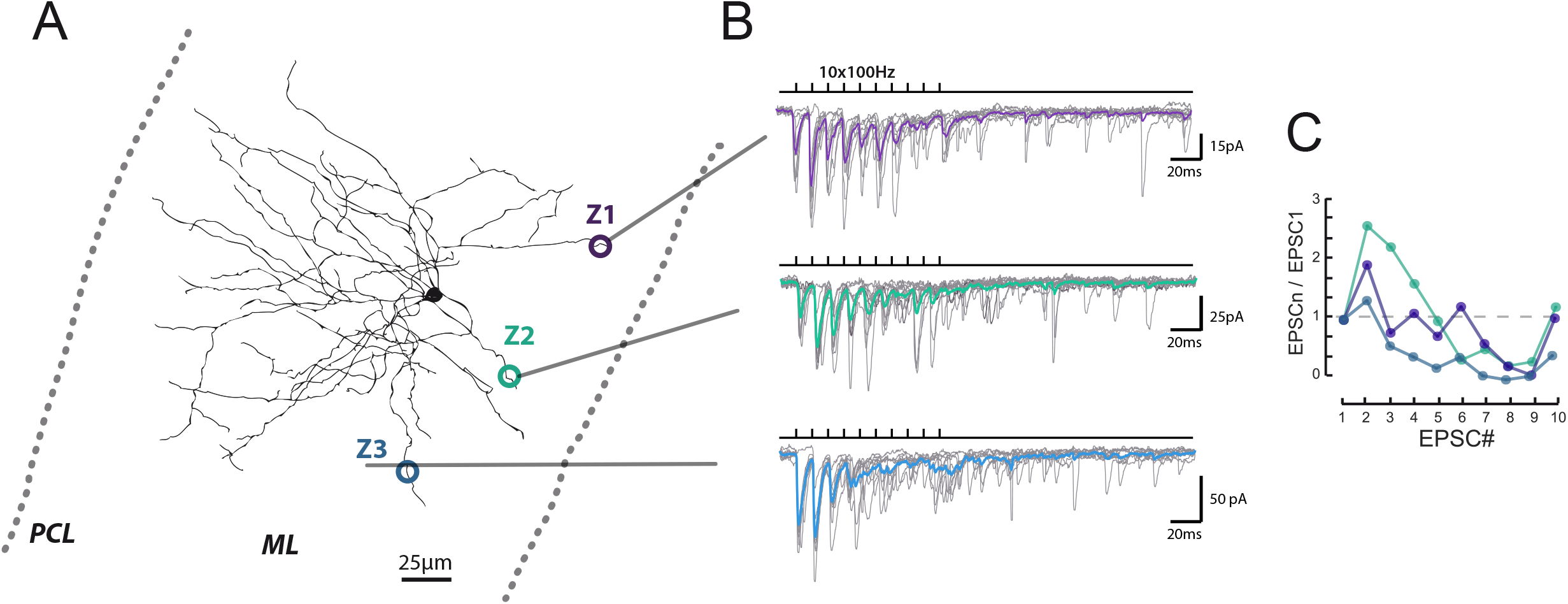
Heterogeneous profile of STP at unitary GC-MLI synapses. (A) Typical experiment showing the profile of STP during 100 trains at 3 unitary inputs recruited by local stimulation of PF at 2 different locations (Z1 to Z3). Figure shows *post-hoc* reconstruction of a recorded MLI. The left and right dashed lines represent the location of the Purkinje cell layer (PCL) and the pia respectively. (B) Superimposed traces correspond to EPSCs recorded during 100 Hz trains after minimal stimulation at Z1, Z2 and Z3 locations. Averaged traces from 10 successive stimulations are represented in purple (Z1), green (Z2) and blue (Z3). (C) Corresponding EPSC charges versus stimulus number at Z1, Z2 and Z3 locations.

**Figure 2.**
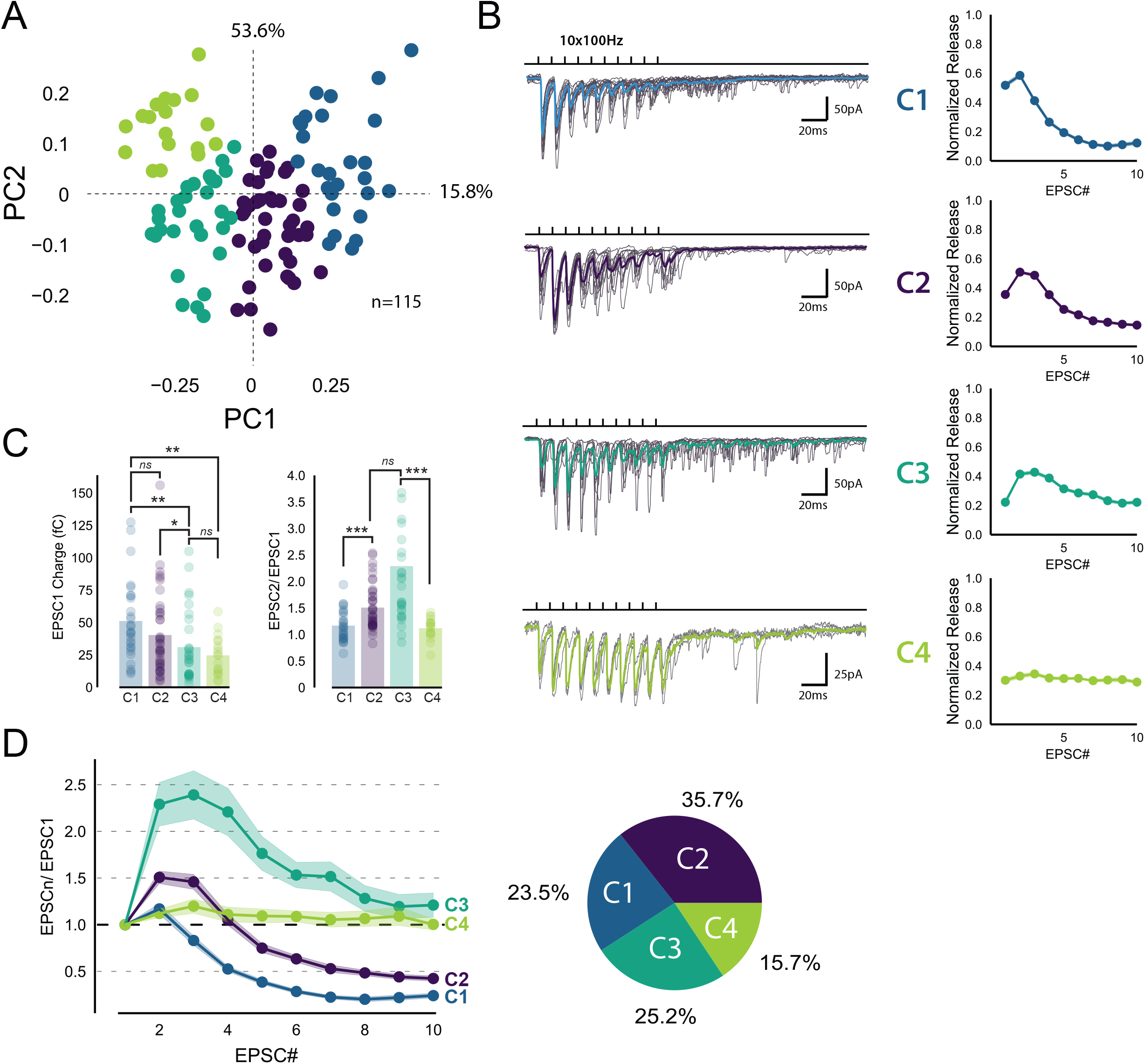
Identification of 4 classes of MC-MLI synapses using PCA followed by fc-means clustering analysis of EPSC properties during high frequency stimulation. (A) PCA transformation of GC-MLI STP profiles. Scatter plot of the first two principal components (PC1, PC2) obtained by analyzing EPSC properties during 100 Hz trains at numerous unitary GC-MLI synapses (*n* = 115). The first two components explain 69.4% of the total variance of STP. Synapses with negative PC1 values sustain glutamate release during the 10 EPSCs of the burst while PC1 with positive values synapses are depressing synapses. Positive PC2 synapses are depressing synapses while negative PC2 synapses are facilitating during EPSC_2,3_ (See supplementary figure 3). (B) Representative traces of the four classes of inputs (C1 to C4) determined by *k*-means clustering analysis during ten minimal stimulation of unitary inputs at 100 Hz. The corresponding values of averaged EPSC amplitude plotted versus the stimulus number and normalized again the vector space model (see method) were displayed on the right panels. (C) *Left panel*, Summary plot of the charge of EPSC recorded at the first stimulus in 100 Hz train (EPSC_1_). (EPSC charge: C1 = 51.29 fC ± 5.96 fC, C2 = 41.63 fC ± 4.97 fC, C3 = 28.80 fC ± 4.78 fC, C4 = 24.75 fC ± 3.54 fC, ANOVA). *Right panel*, Summary plot of the paired-pulse ratio according to the four categories of inputs. (D) Mean values of normalized EPSC amplitudes during 100 Hz train according to the four categories of inputs. The circular diagram represents the relative proportion of each category of input from 96 unitary GC-MLI synapses.

Among the 4 groups identified, only C1 connections exhibited depression of glutamate release after the second pulse, while C2 and C3 connections exhibited facilitation (***Figure 2D*).** C2 and C3 connections differed in their responses after the fourth stimulus: while C3 connections sustained release during the entire train, synaptic transmission at C2 connections depressed after the fourth stimuli. On the other hand, C4 connections were characterized by small but stable EPSCs (***Figure 2C***) suggesting that they correspond to boutons releasing one quanta per stimulation ^30,34^ BCs and SCs receive all profiles of inputs. The absence of cell-specific differences in the first or second principal components (PC1 and PC2) confirms the heterogeneity of STP regardless of MLI subtype (***Figure 3***). Furthermore, no correlation between the first two dimensions of the PCA regarding neither the position of MLI somata nor the localization of synaptic bouton (***Supplementary figure 4***). Thus, the four classes of inputs are randomly distributed within the molecular layer, suggesting that GC inputs contacting a given MLI are heterogeneous.

**Figure 3.**
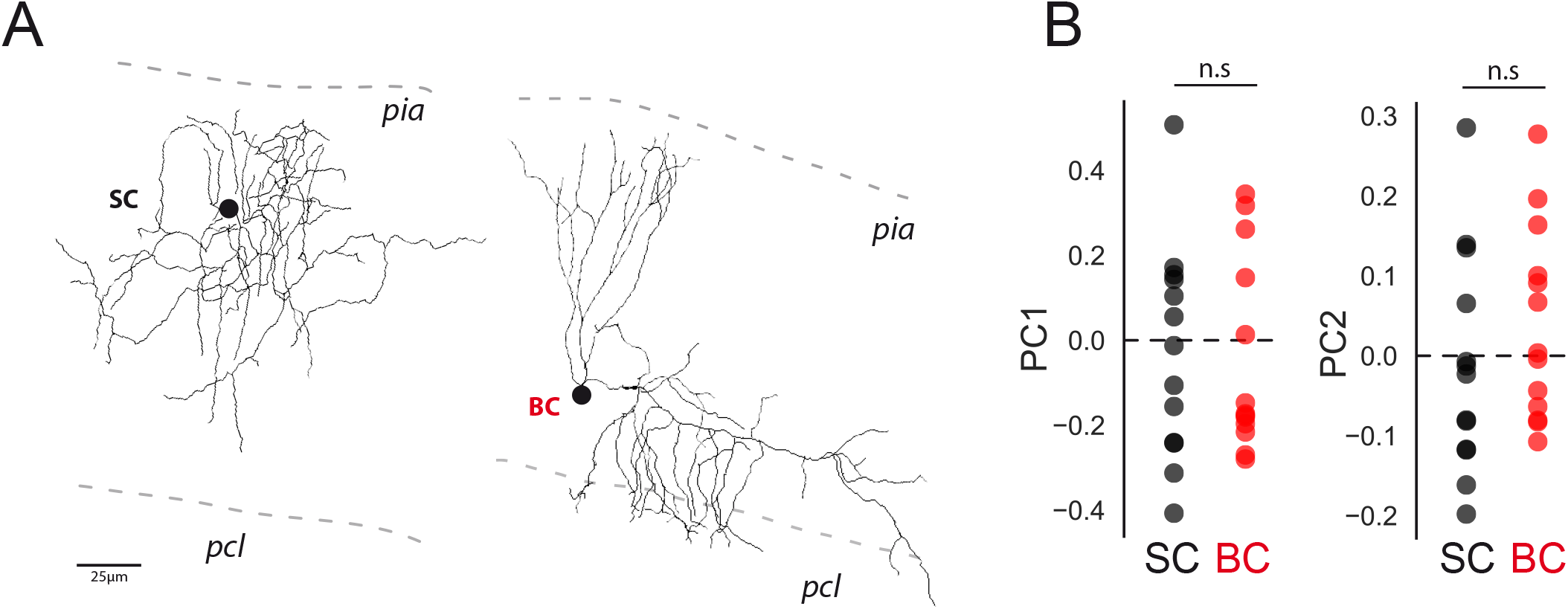
The profile of STP is not determined by the target cell or the position of inputs in the molecular layer. (A). *Post-hoc* reconstruction of 2 recorded MLI using a two-photon microscope. SCs were identified by the absence of neuronal process reaching the PCL (left MLI) and by the absence of cut processes (transection of neuronal processes could be clearly identified by swelling at the tip end portion of processes). At the opposite, BCs were identified by the presence of processes entering in the PCL (right MLI). (B) Scatter plot of PC1 and PC2 obtained at GC-BC synapses (red points) or SC (black points) after minimal stimulation at 100 Hz.

### Syn II is heterogeneously expressed across GC-MLI presynaptic terminals

Next, we aimed to uncover the molecular determinants of the functional heterogeneity. Among presynaptic proteins involved in STP, synapsins (Syn) appeared as good candidates. Syn are presynaptic phosphoproteins coded by 3 distinct genes (Syn I, II and III). While Syn III are mainly expressed during developmental stages, Syn I and Syn II both regulate neurotransmitter release and STP in mature synapses ^31,32,35^ The synapse specific expression of Syn isoforms ^36–38^ contributes to the diversity of STP profiles ^39–42^ and determines the inhibitory-excitatory balance in neuronal networks ^43,44^.

We first studied the presence of Syn I and Syn II in GCs boutons by immunohistochemistry using VGlut1 as specific marker of GC presynaptic terminals ^45,46^. Triple staining of cerebellar sections from P20~P22 CD1 mice (*N* = 4) revealed systematic overlap of VGluT1 with Syn I, but not with Syn II (***Figure 4A*).** Our quantitative analysis revealed a significant correlation between the fluorescence intensity of Syn I and VGlut1, but not between the fluorescence intensity of VGlut1 and Syn II (R_Syn I_ =0.584 +/-0.057, R_Syn II_ = 0.435 +/-0.075, paired t-test: p<0.001; n=52) (***Figure 4A, right panels*).** Our results suggest that Syn I is present in all GC terminals while Syn II is restricted to a subpopulation of GC boutons.

**Figure 4.**
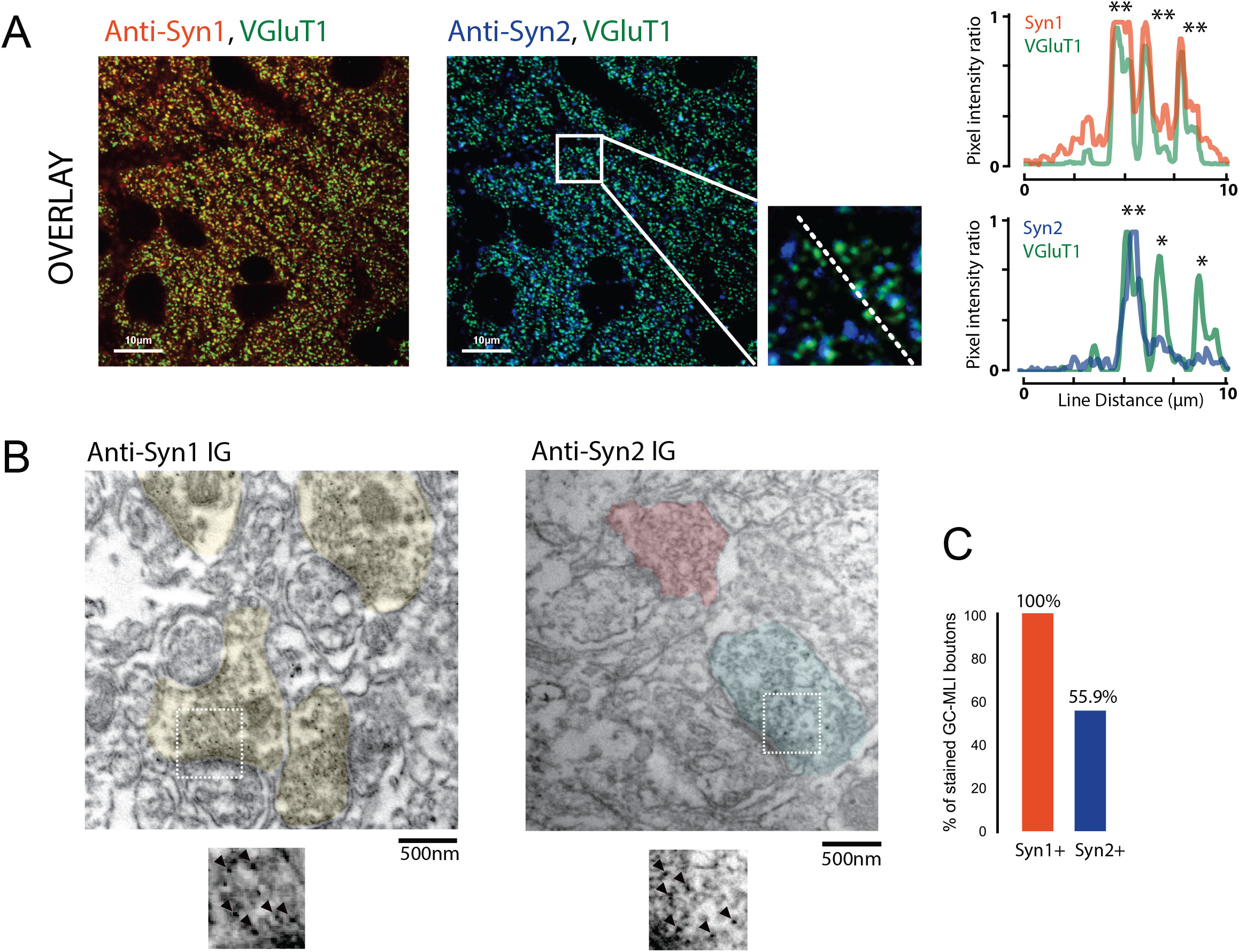
Heterogeneous expression of Syn II at GC-MLI synapses. (A) Representative merged images of VGluT1/Syn I immunostaining (green and red puncta respectively) (left image) or VGluT1/Syn II immunostaining (green and blue puncta respectively) (right image). The two merged imaged were captured in the molecular layer from the same parasagittal cerebellar section. The profile plot (right panel) show the colocalization of VGluT1 with Syn I in the majority of VGluT1 puncta while there was only a partial colocalization of VGluT1 with Syn II in VGluT1 puncta. (B) Typical immunogold electron micrograph illustrating the ubiquitous expression of Syn I in GC boutons contacting MLI (left micrograph) and the heterogeneous expression of Syn II in these boutons (right micrograph). GC boutons contacting MLUI were colorized. Insets corresponding to magnifications of areas delimited by white squares show details of immunogold staining. (C) Histogram of the percentage of GC-MLI synapse positive for Syn I (red bar) and Syn II (blue bar).

To further validate the heterogeneous expression of Syn II in GC-MLI presynaptic terminals, we performed pre-embedding immunogold labeling (***Figure 4B***) of Syn I and Syn II in parasagittal cerebellar sections. Asymmetrical GC-MLI synapses in the upper part of the molecular layer were identified by the presence of mitochondria within the postsynaptic compartment ^26^. Immunogold labeling confirmed the ubiquitous presence of Syn I in all GC-MLI and presence of Syn II in only 56% of GC terminals contacting MLI (***Figure 4C***).

### Heterogeneous expression of Syn II generates diversities in the profile of STP at unitary GC-MLI synapses

The heterogeneous expression of Syn II in GC terminals may contribute to the diversity of STP profiles across unitary GC-MLI synapses. To test this possibility, we reinvestigated STP diversity at unitary GC-MLI synapses in Syn II KO mice (***Figure 5A,B*).** Absence of Syn II modified the responses of unitary GC-MLI synapses to 100 Hz stimulations. The mean EPSC charges of the first responses was strongly reduced in Syn II KO mice and paired-pulse ratio (PPR) was increased (***Figure 5B,C*).** The percentage of failures at the first stimuli were increase in Syn II KO mice indicating that absence of Syn II decreased the probability of release (*p_r_*) of fully-releasable synaptic vesicles (***Figure 5D*).** We then analyzed STP profiles at unitary connections in Syn II-KO as in Figure 2 and plotted individual profiles against the first two dimensions of a PCA based on the PCA fit of WT data (Materials and Methods) and examined the spread of Syn II-KO individual GC-MLI STP observations (***Figure 5E***). The four profiles of STP found at unitary GC-MLI synapses in WT mice were also found at unitary GC-MLI synapses in Syn II KO mice. However, the distribution of the four classes was strongly skewed toward C3 and C4 profiles (85.8% of the connections, n = 33) in Syn II KO mice (***Figure 5 E, F***) while C1 and C2 connections almost disappeared (C1 connection 3.6%, C2 connections 10.7 %) (***Figure 5E***). These results suggest Syn II lead to STP profiles corresponding to C1 and C2, whereas GC boutons displaying C3 and C4 profiles are devoid of Syn II.

**Figure 5.**
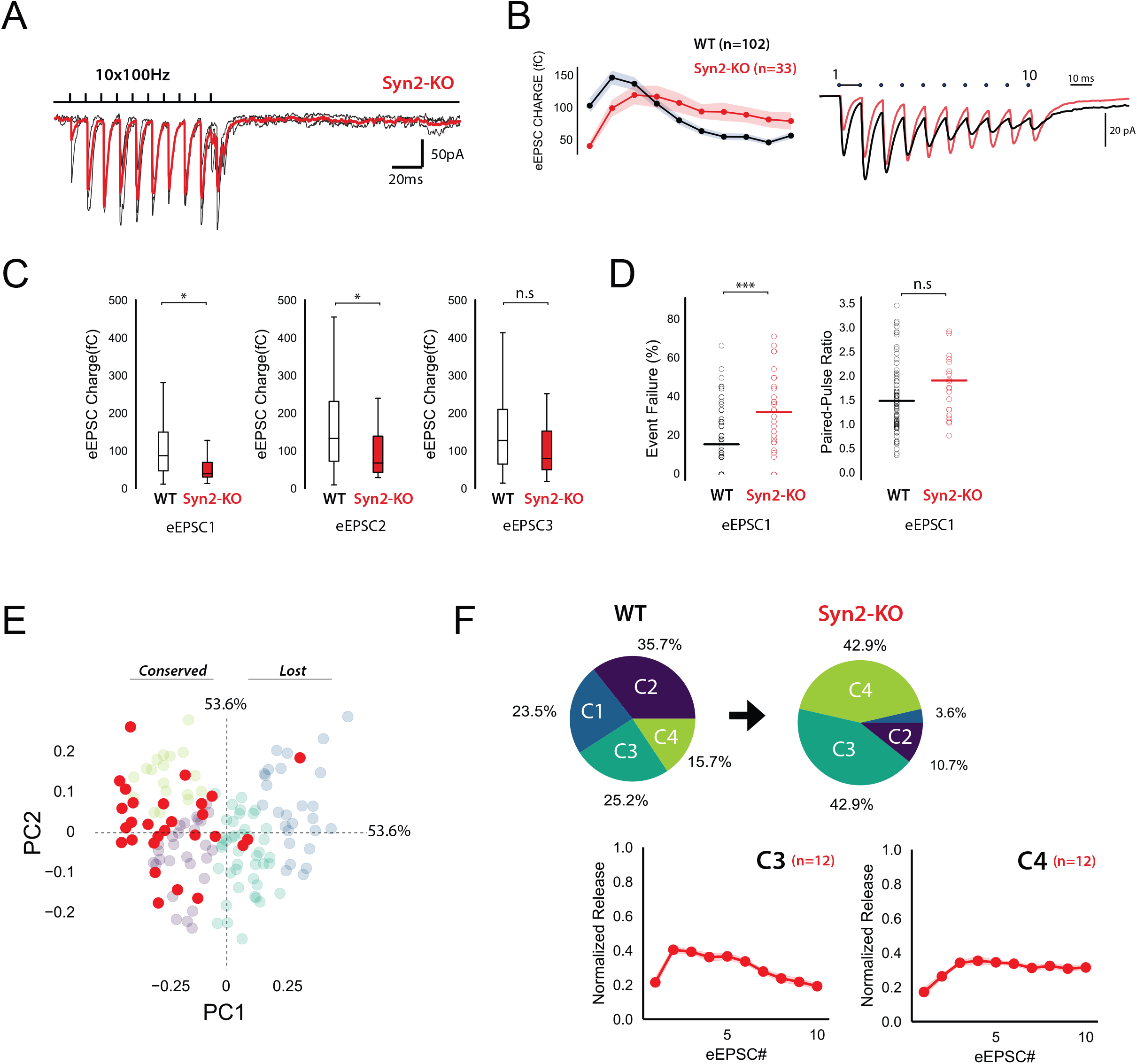
Genetic deletion of Syn II induces a partial loss of functional variability at GC-MLI synapses. (A) Representative traces of averaged EPSCs from 10 successive trains at 100 Hz recorded at unitary GC-MLI synapse from WT (black trace) and Syn II KO mice (red trace). Unitary synapses were stimulated using minimal electrical stimulation. (B) *left panel*, Mean values of EPSC charges elicited by train of stimulation at 100 Hz recorded in WT and Syn II KO mice (black and red points respectively). *Right panel*, Corresponding traces recorded during these 100 Hz train in WT mice (black trace, mean trace from 102 recordings) and in Syn II KO mice (red trace, averaging from 33 recordings). The mean EPSC charges of the first responses was strongly reduced in Syn II KO mice (mean EPSC1 charge: 102.6 fC ± 11.5 fC at WT unitary inputs, *n* = 102, 40 fC ± 7.4 fC, *n* = 33 at Syn KO unitary inputs). (C) Box plots showing the values of EPSC charges at the first, second and third stimulus of 100 Hz train (left, middle and right graph respectively) in WT and Syn II KO mice. (D) Box plots showing the number of failure at the first stimulus and the paired-pulse ratio (left and right graph respectively in WT and Syn II KO mice. The percentage of failures at the first stimuli were increase in Syn II KO mice (mean failure rate EPSC1 in WT: 15.6% ± 1.6, Syn II KO mice: 32.3% ± 3.8) (E) Scatter plot of PCA1 and PCA2 obtained by analyzing EPSC properties during 100 Hz train in WT mice (gray point, same dataset as in Figure 2A) and Syn II KO mice (red points). (F) Pie chart of *k*-means clustering analysis clusters obtained in WT animal (same dataset than in Figure 2C) and Syn II KO mice. Note the near complete disappearance of C1 and C2 connections in Syn II KO mice. The profiles of EPSC charges during 100 Hz train for C3 and C4 connections were identical between WT and Syn II KO mice (bottom graphs), indicating that the genetic deletion of Syn II did not impair the functioning of these two classes of GC-MLI synapses.

We next studied the subcellular localization of synaptic vesicles at GC-MLI synapses from WT and Syn II KO mice using transmission electron microscopy (***Figure 6A*).** Morphometric analysis revealed that the absence of Syn II reduced significantly the number of docked synaptic vesicles and the length of the active zone (***Figure 6B***) without affecting the positive correlation between the length of the active zone and the number of docked synaptic (***Figure 6C*).**

**Figure 6.**
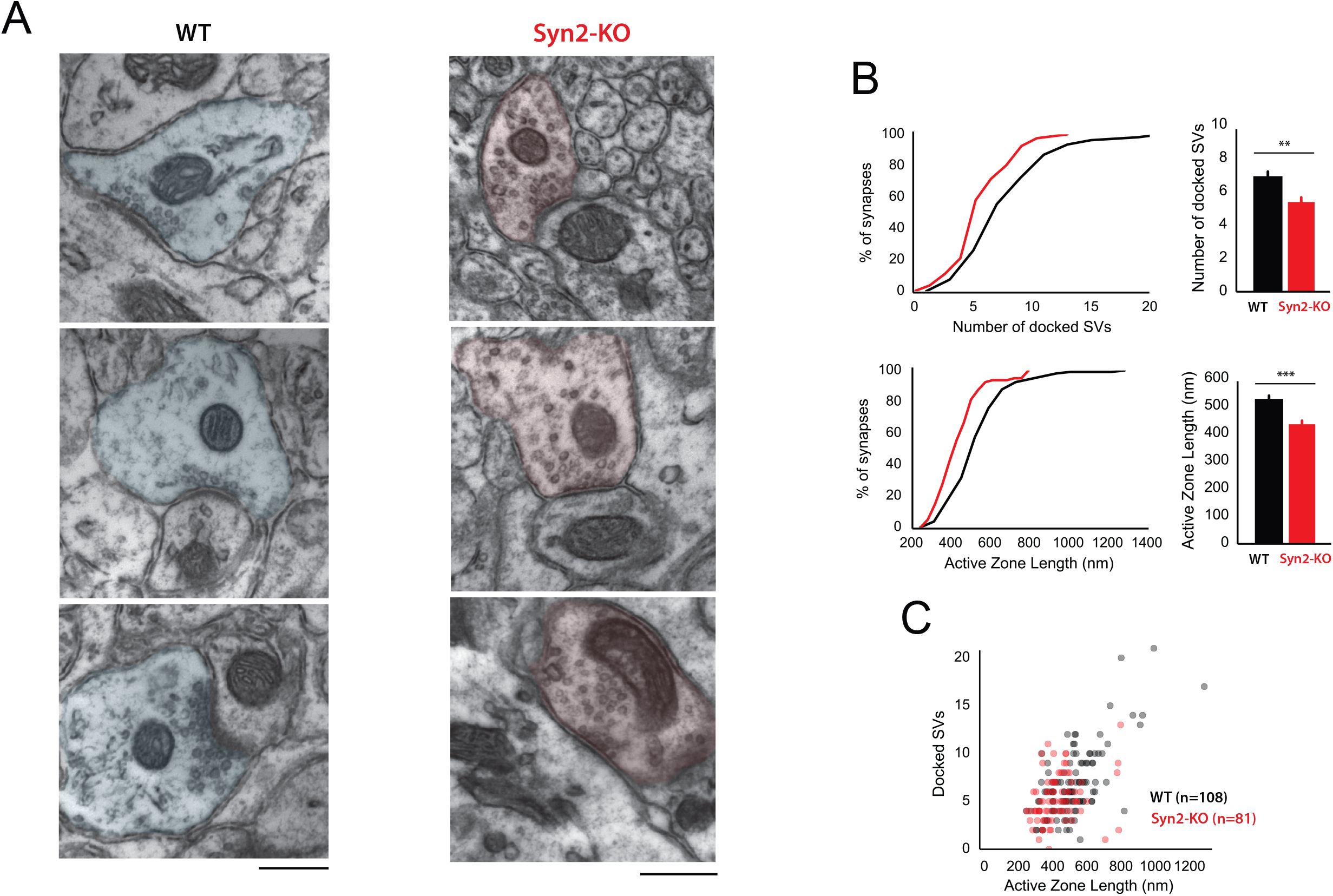
Genetic deletion of Syn II reduces the heterogeneity of ultractructural profiles of presynaptic terminals at GC-MLI. A. Representative micrographs of GC-MLI synapses captured in the upper part of the molecular layer of cerebellar parasagittal sections from WT and Syn II KO mice. (B) *upper panels*, Cumulative distribution (left panel) and mean values of the number of docked SVs at GC-MLI synapses from WT and Syn II KO mice (black line/bar and red line/bar respectively). *Lower panels*, Similar representations for the active zone length. Absence of Syn II reduced significantly the number of docked synaptic vesicles and the length of the active zone (mean number of docked synaptic vesicles: WT 6.83 +/-0.35, n=108; Syn2-KO 5.36 +/-0.28, *n* = 81, mean length of active zone: WT 521.3nm +/-15.2, n=108; Syn2-KO 432.2nm +/-13.2, *n* = 81) (C) Scatter plot of the number of docked synaptic vesicles (SVs) versus ahe active zone (AZ) length from dataset obtained in B. Genetic deletion of Syn II lead to a specific loss of GC bouton endowed with both a long active zone (> 800 nm) and high number of docked synaptic vesicles (> 15 synaptic vesicles).

Altogether, our results suggest that the presence of Syn II positively regulate the number of docked synaptic vesicle and the *p_r_* of fully-releasable vesicles, thus enhancing the release glutamate at the onset of burst firing.

### STP profile controls the temporal coding at GC-MLI connections

The STP profile shapes the spike output pattern of MLIs following compound stimulation of GCs or PFs ^19,47^. This suggests that each class of GC-MLI synapse should influence the MLI spike output pattern specifically. To address this hypothesis, we set out to correlate STP of specific GC units with the spike output pattern of the targeted MLI. We recorded the spike output pattern of MLIs in loose-patch configuration following photostimulation of unitary GC inputs by caged glutamate (Material and Methods and ***Supplementary Figure 5***). Photostimulation of individual GCs increased the MLI firing rate confirming that sufficient glutamate was released by unitary GC boutons during high-frequency stimulation to produce spikes in MLIs ^48,49^ (***Figure 7A*).** Photostimulations, which produced burst in GCs with reproducible parameters (***Supplementary Figure 5***) led to an increase in MLI firing rate (mean baseline frequency: 12.75 ± 5 Hz; peak of acceleration: 33.7 ± 17 Hz, *n* = 32). Subsequently, EPSCs were recorded upon photostimulation of the same unitary GC-MLI synapse using whole-cell configuration (***Figure 7A*).** Photostimulations yielded heterogeneous profile of STP. PCA followed by *k*-means clustering analysis revealed 3 distinct STP profiles (C1’, C2’ and C3’ connections different from C1 to C4 connections since the parameters of bursts elicited in GCs using minimal electrical stimulations and photostimulations are different) that differed by their time course and amplitude (***Figure 7B-C*).** C1’ connections with positive PC1 values were characterized by large responses that peaked at the onset of GC bursts and then rapidly depressed (phasic profile) (averaged EPSC peak charge: −232.8 fC ± 55.2 fC reached at 33.2 ms ± 11.3, *n* = 14). C2’ connections with low PC1 values were also characterized by a phasic profile, but EPSCs have smaller amplitudes (averaged EPSC peak charge: −121.3 fC ± 21.6 fC reached at 59.7 ms ± 10.3, *n* = 30) than C1’ synapses. C3’ connections were characterized by smaller responses that peaked with longer delays than the ones of C1’ or C2’ synapses (EPSC peak charge: −133.4 fC ± 21.9 fC reached at 84.4 ms ± 5.4, *n* = 18).

**Figure 7.**
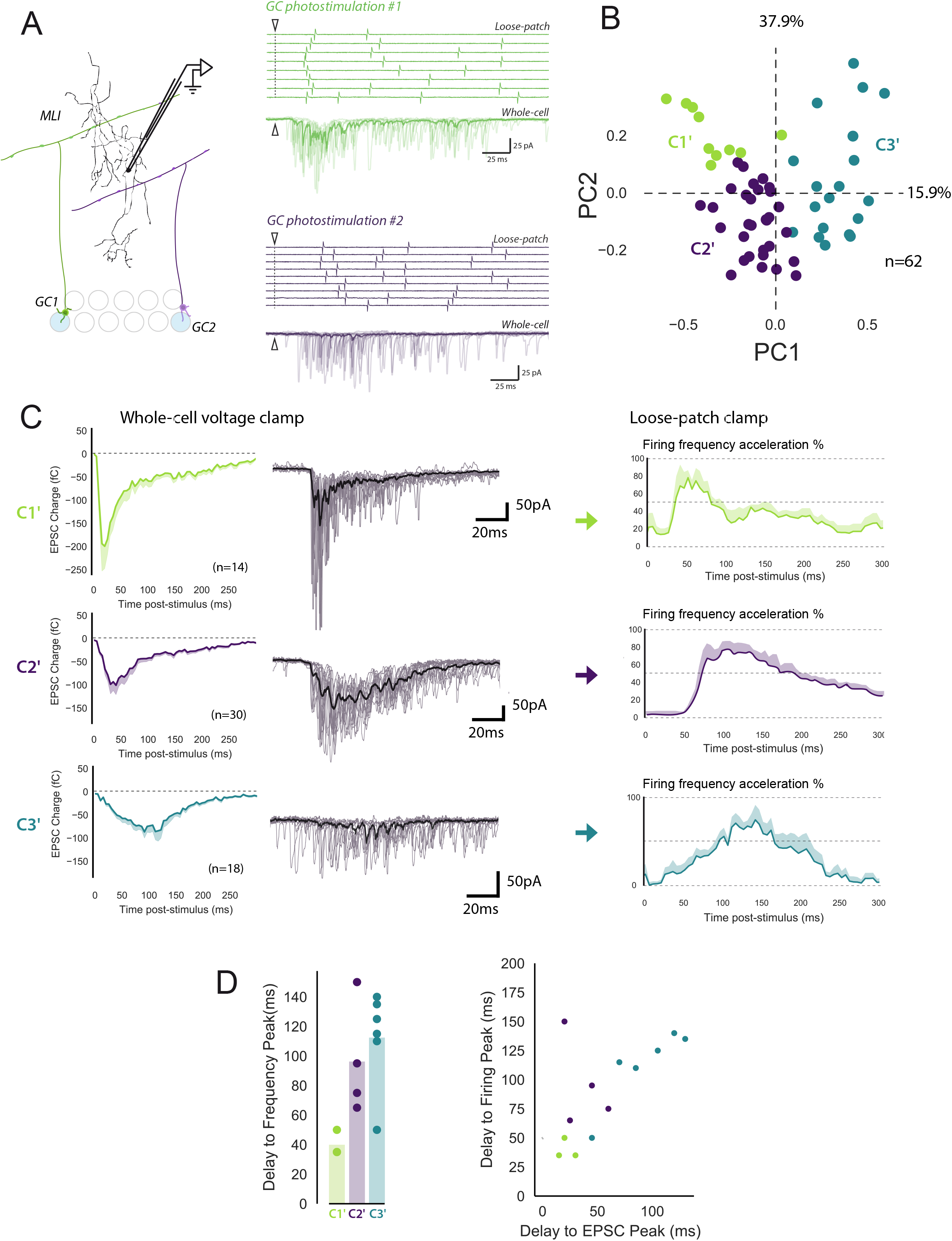
The diverse profiles of STP of GC boutons differentially shape the spike output pattern in the target MLI. (A) *left panel*, Schematic representing the design of photostimulation in the GCL. Open circles represent the sites where RuBi-glutamate was uncaged and the blue circles represent the locations where photostimulation elicited responses in the recorded MLI. In this example, 2 GCs localized at distal positions in the GCL contact the recorded MLI. *Right panels*, Representative experiment showing the spike output pattern recorded in loose-patch configuration and EPSCs recording in whole cell configuration in the same MLI following photostimulation of two different locations in GCL. The white arrowheads and dashed lines represent the onset of photostimulation. Note that the onset of firing is time-locked to the first peak of EPSC charge for photostimulations in location #1 (upper panels) while the onset of firing was more variable for photostimulation in location #2 (lower panels). (B) PCA transformation of the evoked charge time course for 63 unitary contacts (see methods). The EPSC bursts could be differentiated depending on their tonic or phasic component into three different clusters using k-Means clustering. (C) Within the population of synaptic terminals, some responded with phasic and depressing EPSC burst (cluster1), others facilitated and still have a lower phasic component (cluster2). The third cluster is characterized by a sustain release of glutamate with low amplitudes. We retrieved loose patch recordings of MLI output firing pattern and we associated synapse-specific MLI firing patterns to GC-MLI STP from the three clusters (*right panel*). We show a stereotypic response of MLI depending on the category of GC-MLI STP that has been activated. (D) Heterogeneous presynaptic boutons with different GC-MLI STP evoke specific activations of MLIs. *Left panel*, bar plot representing the values of the delay to frequency peak for each clusters of synapses classified using *k*-Means clustering. *Right panel*, The scatter plot shows a high correlation between the peaks of EPSCs recorded in voltage-clamp condition and the delay to frequency peak recorded in loose-patch configuration. Each point corresponds to values obtained at unitary GC-MLI synapses. The color code which is identical to the left graph identify the cluster is which each connections were classified.

In 12 unitary connections out of 62, we could correlate the spike output pattern recorded in loose-patch configuration with the STP profile recorded in whole-cell configuration (***Figure 7C right panel*).** Photostimulation of C1’ connections accelerated the firing rate of the targeted MLI with a very short delay (< 50 ms) compared to C2’ and C3’ connections (delay >60 ms). Our analysis also showed a clear relationship between PC1 and the peak frequency (Pearson coefficient, R= 0.7, *p* = 0,008, *n* = 13) (***Figure 7D*).** This indicates that a specific STP profiles determines the delay to the first spike in MLI in response to GC photostimulation. These results suggest that the properties of glutamate release at GC-MLI synapses during high-frequency stimulation are major determinant of the temporal coding of sensorimotor inputs through the FFI pathway.

We next tested how Syn II deficiency affects the correlation between STP profiles ad firing pattern at GC-MLI synapses, using Syn II KO mice (*n* = 27). PCA followed by *k*-means clustering analysis showed that genetic deletion of Syn II almost abolished the presence of C1’ connections and nearly doubled the percentage of C3’ connection (***Figure 8A-D*).** Moreover, the delay to the first spike significantly increased in Syn II KO mice (***Figure 8E,F*),** probably due to a lower initial *p_r_* at GC-MLI synapses devoid of Syn II (***Figure 5 A,B*).** Our results reveal Syn II a major determinant of fast temporal coding at the GC-MLI synapses. Synapse-specific expression of Syn II diversifies the profile of excitatory drives on MLIs and expands the temporal coding in the FFI pathway.

**Figure 8.**
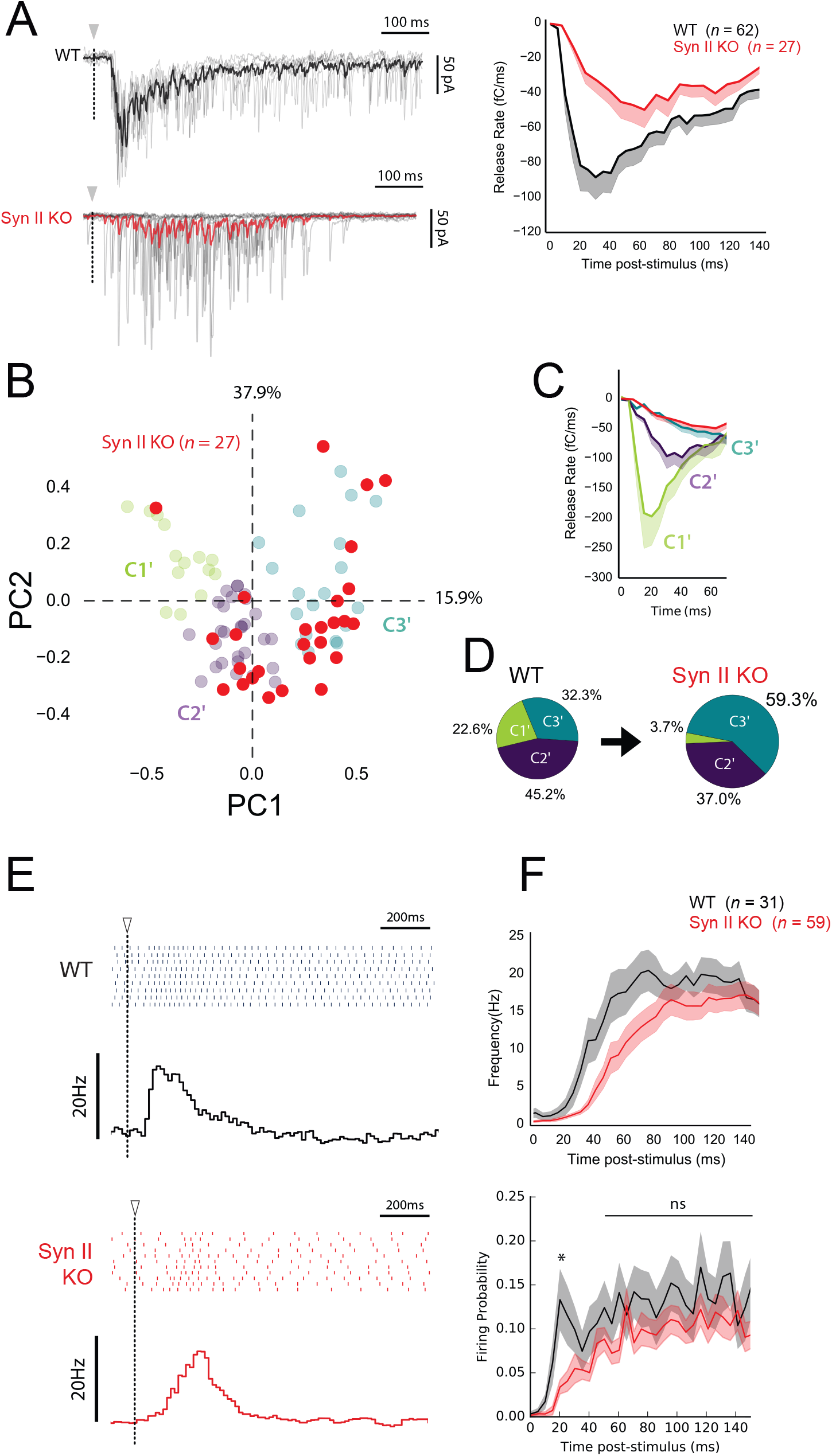
Syn II expands temporal coding at GC-MLI synaptic synapses. (A) *left panels*, Representative traces of superimposed EPSCs recorded in WT and Syn II KO mice after photostimulations in the GCL (same experimental design than in figure 7A). *Right panel*, Averaged values of EPSC charge versus the time following photorelease of RuBi-glutamate recorded in WT and Syn II KO mice (black and red traces respectively). Note the strong reduction in the peak of charge in Syn II KO mice. (B) PCA transformation of EPSC properties obtained in WT (green, purple and blue points, same dataset than in Figure 7B) and Syn II KO mice (red points). (C) Line plots display the normalized release time course from GC-MLI synapses belonging to clusters C1’, C2’, C3’ (WT mice, same color code as in B) and GC-MLI synapse from Syn2-KO mice (red line). (D) The pie chart shows a partial reduction of STP heterogeneity in Syn2-KO condition with a strong reduction of phasic profiles (C1’). (E) Typical raster plots and peristimulus time histogram obtained in WT and Syn II KO mice following photostimulation of unitary GC-MLI synapses. The onsets of photostimulation are represented with white arrowheads and dashed lines. (F) Means values of the firing frequency (upper graph) and the firing probability (lower graph) of MLIs following photostimulation of unitary GC-MLI synapses in WT and Syn II KO mice (black and red lines respectively).

## DISCUSSION

By combining molecular, ultrastructural and functional studies, we show that the firing pattern of MLIs is driven by distinct GC inputs that show distinct profile of STP. This functional heterogeneity caused, at least in part, by synapses-specific expression of Syn II expands the temporal coding of GC inputs to MLIs.

### MLI subtype does not determine the profile of STP at unitary GC-MLI synapse

The rules governing diversity in presynaptic release properties have been extensively studied in neocortical or hippocampal circuits. In most of cases, synaptic efficacy among boutons issued from a single axon varies with the identity of the postsynaptic cell ^50,51^. Target cell-dependent heterogeneity relies on differences in the probability of release ^52^, responsiveness to neuromodulators ^53–56^ or the ability to co-release GABA and glutamate ^57^. Target cell-dependent STP has also been described at cerebellar GC-MLI synapses by using stimulation of beam of PFs or clusters of GC somata; upon high-frequency stimulation, compound synaptic responses exhibit a facilitating profile at GC-SC synapses whereas these responses depress at GC-BC synapses ^19^. Our results argue against target-cell dependent STP at GC synapses and show rather heterogeneous unitary inputs to BC- or SC-like MLIs (***Figure 1,3*).** Several hypotheses may explain the discrepancies between results obtained with compound stimulations versus unitary ones. In compound responses, connections associated with strong synaptic strength mask the influence of weak inputs. Also, target-cell dependency of STP may be reduced to some MLIs localized near the pia (SC-like MLIs) or at the opposite in the direct vicinity of the PC layer (BC-like MLI). Anyhow, our work suggests that the excitatory drive to MLIs and the tuning of the FFI pathway are more complex than expected.

### Organization of synaptic diversity at unitary GC-MLI synapses

Heterogeneous expression and functions of Syn II in different cell types forming a same neural network have been reported previously ^36–38,40–42^ but our findings bring the first evidence that expression of Syn II at a given connection can be heterogeneous. Syn II expression may be genetically determined at early developmental stages, leading to Syn II(+) and Syn II(-) subclones of GCs. It is interesting to note that clonally related GCs (that is, GCs issued from the same GC progenitors) stack their axons in a specific sub-layer in the molecular layer ^58^ suggesting that the presence or absence of Syn II may be organized in a beam-dependent way. Since beams of neighboring PFs are activated during sensory stimulations ^59^, recruitment of Syn II(+) or Syn II(-) connections may be related to the activation of a given sensorimotor task. Alternatively, Syn II targeting at individual PF boutons may be controlled by a complex interplay of mechanisms regulating the traffic of Syn in axons ^40^ or organizing the assembly of the presynaptic active zone^60^.

Presynaptic diversity also arises from other parameters that probably expand the range of synaptic behaviors across GC boutons. Calcium imaging performed on single PFs revealed that Ca^2+^ dynamics and regulation of Ca^2+^ influx by neuromodulators in synaptic varicosities from a same PF are highly heterogeneous ^61–64^ Also, local retrograde release of endocannabinoid by MLI dendrites can affect the functioning of subsets of GC boutons upon sustained activity of PF ^29,65^ Hence, functional heterogeneities among GC terminals also originate from the history of firing of each GC. To summarize, the synaptic behavior of individual GC bouton may be tuned by an intermingled combination of factors including expression of Syn II at GC-MLI synapses, presynaptic receptor composition, presynaptic Ca^2+^ dynamic, number of active release sites, retrograde signaling and history of firing.

### Control of glutamate release by Syn II in GC boutons

At GC synapses, releasable synaptic vesicles are segregated in two pools, one with fully releasable vesicles and a second one with reluctant vesicles, which are differentially poised for exocytosis ^14^. The fully-releasable pool supports glutamate release during single action potentials while the reluctant pool is recruited only by stimuli elicited at high-frequencies ^14^. In Syn II KO mice, synaptic transmission is characterized by a defect in glutamate release by single action potentials and by a rapid recovery of synaptic transmission by 100 Hz stimuli. This suggests that a lack of Syn II impair *p_r_* of fully-releasable vesicles without affecting the recruitment of reluctant vesicles. Potentially, Syn II may act with several partners to control the recruitment of fully-releasable vesicles. In GC terminals, Munc13-3 has been involved in superpriming steps that tightly couple synaptic vesicles with P/Q-type Ca^2+^ calcium channels (positional superpriming) or maturate the fusion machinery (molecular superpriming) ^66–68^. Munc13-3 may indirectly act with Rab3-interacting molecules (RIMs) which are well known organizers of calcium channel and synaptic vesicles in the active zone ^69^. Since Syn II interacts with both Rab3 ^70^ and P/Q type calcium channels ^71^, it cannot be excluded that Munc13-3, Syn II, Rab3 and RIM act in concert to reduce the physical distance between fully-releasable vesicles and Ca^2+^ channels. Alternatively, Syn II-Rab3-RIM complex may directly regulate the influx of Ca^2+^ through strong inhibition of voltage-dependent inactivation of P/Q type Ca^2+^-channels ^72,73^.

### Physiological consequences

At the input stage of the cerebellar cortex, single GCs receive a combination of MF inputs coding for different modalities ^74,75^. The diversity of STP profiles across MFs from different origins provide temporal signatures for each combination of MFs converging on a single GC thus enhancing pattern decorrelation of sensory inputs ^76^ Here we show that temporal coding in GCs is later extended in the FFI pathway by an input-specific control of first-spike latency in MLIs. The combination of heterogeneous presynaptic behaviors at the successive stages of cerebellar computation leading to consecutive temporal signatures, should refine the salient feature of a given combination of MF inputs and ultimately should enhances the representation of sensory information by PCs.

Considering the importance of delay coding for internal models of motor adjustments ^77–79^ synapse-specific temporal coding may have essential consequences for learning and predictive functions in the cerebellum. Indeed, longterm potentiation (LTP) of GC-PC connections require the coincidence of strong PF and MLI activities onto the same PC ^80^. As exemplified by the rules governing the induction of associative GC-PC long-term depression triggered by coincident PF and climbing fiber activation ^81^, induction of associative GC-MLI LTP may depend on the time window separating excitatory and inhibitory inputs onto PCs. Heterogeneous profiles of STP at MF-GC synapses and GC-MLI synapses necessarily induce wide range of delays between the direct excitatory pathway and FFI at the level at PC synapses. Hence, the fine tuning of STP at the level of single cells may have fundamental importance for the induction of long-term plasticity and ultimately for motor learning.

## Material and Methods

This study was carried out in strict accordance with the national and international laws for laboratory animal welfare and experimentation and was approved in advance by the Ethics Committee of Strasbourg (CREMEAS; CEEA35; agreement number/reference protocol: APAFIS#4354-20 16030212155187 v3). Mice were bred and housed in a 12 h light/dark cycle with free access to food and water. Wild type (WT) or Synapsin II knock-out (Syn II KO) mice have CD1 genetic background. Syn II KO mice were first derived from synapsin triple knock-out mice (C57BL/6J genetic background, originating from the Italian Institute of Technology, Genova, Italy) ^40^ bred with CD1 WT mice. Syn II KO hybrid mice were serially bred (10 backcrosses) with CD1 WT mice to obtain Syn II KO mice with CD1 genetic background.

### Slice preparation

Acute cerebellar slices were prepared from CD1 mice or Syn II KO mice ^82^, aged 20 to 35 days. Syn II KO mice. Mice were anesthetized by isoflurane ihhalation and decapitated. The cerebellum was extracted in ice-cold (~1°C) artificial cerebrospinal fluid (ACSF) bubbled with carbogen (95% O_2_, 5% CO_2_) containing (in mM): 120 NaCl, 3 KCl, 26 NaHCO3, 1.25 NaH_2_PO_4_, 2.5 CaCl_2_, 2 MgCl_2_, 10 D-glucose and 0.05 mM minocyclin. Cerebella were sliced (Microm HM650V, Germany) in an ice-cold low-sodium and zero-calcium slicing buffer containing (in mM): 93 N-Methyl-D-Glucamine, 2.5 KCl, 0.5 CaCl2, 10 MgSO4, 1.2 NaH2PO4, 30 NaHCO3, 20 HEPES, 3 Na-Pyruvate, 2 Thiourea, 5 Na-ascorbate, 25 D-glucose and 1 mM Kynurenic acid. Sagittal or horizontal slices 300 μm thick were immediately transferred for recovery in a bubbled ACSF for 30 minutes at 34°C and maintained at room temperature (~25°C) in bubbled ACSF before use.

### Electrophysiology

After at least 1 hour of recovery at room temperature (~25°C), slices were transferred in a recording chamber continuously perfused with 32~34°C bubbled ACSF. In order to block all forms of long term synaptic plasticity and trans-synaptic signalling, blockers of GABA_A_-receptors (100μM picrotoxin), GABA_B_-receptors 10 μm (3-[[(3,4-Dichlorophenyl)-methyl]amino]propyl(diethoxymethyl)phosphinic acid), NMDA-receptor (100 μM D-AP5; D-(-)-2-Amino-5-phosphonopentanoic acid), endocannabinoïd CB1 receptors (1 μM AM251 1-(2,4-Dichlorophenyl)-5-(4-iodophenyl)-4-methyl-N-(piperidin-1-yl)-1H-pyrazole-3-carboxamide) and mGluR1 receptor (2 μM JNJ16259685 (3,4-Dihydro-2H-pyrano[2,3-b]quinolin-7-yl)-(cis-4-methoxycyclohexyl)-methanone) were added in ACSF.

MLI were patch-clamped in lobules IV to VI in the cerebellar vermis using a two-photon microscope setup (Multiphoton Imaging System, Scientifica UK) with 10 MΩ resistance glass electrodes containing a cesium-based intra-cellular medium (140 mM CsCH3SO3, 10 mM Phosphocreatine, 10 mM HEPES, 10 mM BAPTA, 4 mM Na-ATP and 0.3 mM Na-GTP) supplemented with 50 μM ATTO-594 fluorescent dye. In all experiments, cells were voltage-clamped at −70mV in whole-cell configuration (Multiclamp 700B, Molecular Devices). Data were acquired using the WinWCP freeware (John Dempster, Strathclyde Institute of Pharmacy and Biomedical Sciences, University of Strathclyde, UK). Electrical stimulations were realized with a ~10 MΩ resistance monopolar electrode also filled with ATTO-594 for a precise adjustment of the distance between the stimulation pipette and isolated dendritic processes of MLIs. Electric pulses were adjusted at any GC-MLI contact and evoked with a stimulator (IsoStim01-D, NPI Germany)

Minimal stimulation was used to monitor short term plasticity at unitary GC-MLI synapses in sagittal slices. To do so, we followed previously established procedures For each synapse, the intensity of electrical stimulation was maintained in an intensity window that avoided both stimulation failures and multiple-synapse stimulation (***Supplementary figure 1***). STP at GC-MLI synapses was assessed with 10 trains of ten pulses at 100Hz train separated by a resting period of 1 minute.

We performed glutamate-uncaging assays onto horizontal slices by using MOSAiC patterned illumination system (Andor Technologies). MLI were recorded in ACSF containing 100 μM RuBiGlutamate ^83^ In order to find connected pairs of GC-MLI, we first used full-field arrays composed of very small photostimulation areas (15~25μm diameter) and patches of GC were sequentially illuminated with blue light (460 nm). We took advantage on horizontal slice configuration to stimulate GCs localized at distant locations from the recorded MLI. Considering the weak probability of connection between GC and MLI, synaptic activities evoked by photostimulation of small cluster of GCs localized far away of the recorded MLI are likely to originate from unitary GC-MLI synapses.

### Post-hoc 3D reconstructions

After the experiments, two-photon Z-stacks (1 μm resolution) were done to reconstruct the recorded MLIs in sagittal configuration using the *simple neurite tracer* plugin ^84^ from ImageJ freeware (National Institute of Health, USA). Basket cells were identified by the basket-like features observed in the Purkinje cell layer ^26^.

### Electron microscopy

CD1 mice aged 20 days (WT and Synapsin 2 KO mice) were deeply anaesthetized by intra-peritoneal injection of Ketamine (2 ml/kg) and Xylazine (0.5 ml/kg) and intracardiac perfusion was performed with 2,5% glutaraldehyde in phosphate buffer (0.1 M, pH 7.4). For immunogold labeling, the fixative solution was replaced by 0,1% glutaraldehyde and 4% paraformaldehyde in phosphate buffer. Transversal cerebellar vibratome sections (75 μm thick) were cut and processed either for ultrastructural analysis or for pre-embedding immunogold labeling. After three washes in phosphate buffer, sections were post-fixed in phosphate buffer with 1% OsO4 for 1 hour. Slices were dehydrated in a graded alcohol series (ethanol 25 %, 50 %, 70 %, 95 % 100 %; 10 min per bath) except for ethanol 100 % (3×10 min) followed by an incubation in propylene oxide for 3×10 min. Then slices were embedded in Araldite M (wash in propylene oxide at 1:1 for 1 hours followed by Araldite M for 2×2 hours at room temperature; polymerization at 60°C for 3 days). Ultrathin sections were finally contrasted with uranyl acetate before.

### Pre-embedding immunogold labelling

Sections were permeabilised with 0.2% saponin in PBS, rinsed in PBS and blocked in a blocking solution: 2% bovine serum albumin in PBS (PBS-BSA). The sections were incubated overnight with anti-Syn1 (1/250) or anti-Syn2 (1/100) antibodies (polyclonal, SynapticSystems) in 0.1% BSA in PBS. After washing in PBS-BSA, the sections were incubated in Ultra small nanogold F(ab’) fragments of goat anti-rabbit or goat anti-mouse immunoglobulin G (IgG) (H and L chains; Aurion) diluted 1/100 in PBS-BSA. After several rinses in PBS-BSA and in PB, sections were postfixed in glutaraldehyde 2% in PB before washing in PB and distilled water. Gold particles were then silver enhanced using the R-Gent SE-EM kit (Aurion) before being washed in distilled water and PB. Finally, the sections were post-fixed in 0.5% OsO4 in PB for 10 min before classical processing for Araldite embedding (Sigma, St. Louis, MO, USA) and ultramicrotomy. The ultrathin sections were counterstained with uranyl acetate and observed with a Hitachi 7500 transmission electron microscope (Hitachi High Technologies Corporation, Tokyo, Japan) equipped with an AMT Hamamatsu digital camera (Hamamatsu Photonics, Hamamatsu City, Japan). In control sections processed without anti-Syn1 or anti-Syn2 primary antibodies or gold-labeled secondary antibodies, no gold particles were observed.

### Analysis of electron micrographs

GC-MLI synapses were recognized by the following criteria: i) these synapses are asymmetrical ^85^ ii) presence of mitochondrion within the postsynaptic compartment^86^. Morphometric analyses were performed using ImageJ freeware (National Institute of Health, USA). Then we binned the number of vesicles (with 50nm distance bins) starting from the active zone cytomatrix as a reference point (0nm).

### Immunohistochemistry

CD1 WT mice aged 20 to 25 days were deeply anaesthetized by intra-peritoneal injection of Ketamine (2 ml/kg) and Xylazine (0.5 ml/kg) and perfused with phosphate buffer saline (PBS) containing 4% paraformaldehyde (PFA). After a 3 hour post-fixation, cerebella were sliced in sagittal configuration (50μm thickness). Slices were washed in PBS (3×10 minutes). Membranes were permeabilized by 0,1% TritonX100 and non specific antigens were blocked by 10% bovine serum albumin (BSA) and 1% goat serum albumin (GSA) during 6 hours. Synapses were stained using the same solution supplemented with anti-VGluT1 polyclonal antibodies diluted at 1/600 (SynapticSystems) and polyclonal pan-Synapsin I and pan-Synapsin II antibodies (SynapticSystems) diluted at 1/500. Secondary antibodies (Abcam) were applied during 3 hours in a solution containing 10%BSA. Slices were mounted and visualized under confocal microscope (Leica SP5, II).

### Data analysis

Analysis were performed with home-made python routines (WinPython 3.3.5, Python Software Fundation) based on custom scripts. All statistical analysis were performed using SciPy plugin (https://scipy.org/). Error bars represent +/-SEMs of data distribution. Student’s t-test was used in the case of a normal distribution of data, Mann-Whitney Rank Sum Test (MWRST) was used in other cases. The levels of significance are indicated as ns (not significant) when p>0.05, * when p 0.05, ** when p < 0.01 and *** when p < 0.001.

### Principal Component Analysis (PCA)

PCA is a linear transformation algorithm that examines the main sources of variability inside a dataset composed of multiple observations in order to classify the dataset. PCA analyses covariance between the n variables of a dataset and transforms an original dataset in *eigenvalues* around a small number of dimensions representing the principal components. The first two Principal Components (PC1 and PC2) which explain the highest source of variance from the original dataset are represented in a scatter plot. We used PCA in order to classify STP in our datasets and extract the most relevant inter-individual differences. PCA were computed using the python-based *sklearn* plugin. Input variables were normalized and centered using Vector Space Model (VSM) that linearly scales the observations between 0 and 1 (Salton, Wong, & Yang, 1975). While STP data from WT GC-MLI terminals was used for PCA computation, Syn II KO observations did not take part in the eigenvalue calculation. In order to compare STP heterogeneity between the two populations of synapses, Syn II KO observations were processed as additional values and overlaid to WT cloud of points

### Data processing

For STP analyses using minimal stimulation protocols, data was collected by estimating EPSC charges at any stimulus number from 7 (or more) consecutive trains at 100 Hz elicited every minutes. Failures were arbitrarily detected as signals below a threshold of 3 × σ_noise_, where σ_noise_ is the standard deviation of the amplitude of the noise calculated on a 300ms fixed temporal window preceding the stimulation. PCA transformations (***Figure 2,5,7***) were performed on the median charge value of each EPSC from the 100Hz train pulse for each synapse (n=96). The charge of EPSCs evoked at unitary GC-MLI synapses by photostimulation was measured in a minimal number of 7 successive recordings. To calculate the average charge, values were binned (bin width = 5ms) from the stimulation onset to 100 ms post-stimulus for each sweep (n=1080) and PCA transformation was applied using the charge value for unitary dataset (***Figure 6,7 n=89*).** The delay of MLI peak frequency was estimated from the stimulation onset.

In rare cases, photostimulating GCs could evoke neurotransmitter release at more than one GC-MLI synaptic contact. To optimize GC-MLI STP dataset, *post-hoc* monitoring of EPSCs evoked by GC activation was systematically performed with Igor PRO (WaveMetrics), using the SpAcAn plugin http://www.spacan.net/. When EPSCs displayed important kinetic variability, the recordings (both loose-patch and whole-cell recordings) were systematically discarded.

**AUTHOR CONTRIBUTIONS**
FD, KD and PI designed the study. KD performed electrophysiological and immunohistochemical experiments. VD performed electron microscopy experiments. BP, YB provided theoretical and analytical tools for analyses. KD analyzed data. FD, KD and PI wrote the manuscript.

## ACKNOWLEDGMENTS

This work was supported by the Centre National pour la Recherche Scientifique, the Université de Strasbourg, the Agence Nationale pour la Recherche Grant (ANR-2015CeMod) and by the Fondation pour la Recherche Médicale to PI (# DEQ20140329514). KD was funded by a fellowship from the Ministère de la Recherche.

We thank Sophie Reibel-Foisset and the staff of the animal facility (Chronobiotron, UMS 3415 CNRS and Strasbourg University) for technical assistance. We thank Pr. Fabio Benfenati (Italian Institute of Technology, University of Genova, Genova, Italy) for the gift of synapsin triple knock-out mice.

## REFERENCES

1. Hennequin, G., Agnes, E. J. & Vogels, T. P. Inhibitory Plasticity: Balance, Control, and Codependence. Annu. Rev. Neurosci. 40, 557–579 (2017).

2. Klausberger, T. & Somogyi, P. Neuronal Diversity and Temporal Dynamics: The Unity of Hippocampal Circuit Operations. Science (80-.). 321, 53–57 (2008).

3. O’Donnell, B. V., Tew, D. G., Jones, O. T. & England, P. J. Studies on the inhibitory mechanism of iodonium compounds with special reference to neutrophil NADPH oxidase. Biochem. J. 290 (Pt 1, 41–9 (1993).

4. Isaacson, J. S. & Scanziani, M. How inhibition shapes cortical activity. Neuron 72, 231–243 (2011).

5. Jörntell, H., Bengtsson, F., Schonewille, M. & De Zeeuw, C. I. Cerebellar molecular layer interneurons - computational properties and roles in learning. Trends Neurosci. 33, 524–32 (2010).

6. Jelitai, M., Puggioni, P., Ishikawa, T., Rinaldi, A. & Duguid, I. Dendritic excitation–inhibition balance shapes cerebellar output during motor behaviour. Nat. Commun. 7, 13722 (2016).

7. Armstrong, D. M. & Edgley, S. a. Discharges of interpositus and Purkinje cells of the cat cerebellum during locomotion under different conditions. J. Physiol. 400, 425–445 (1988).

8. Ozden, I., Dombeck, D. a., Hoogland, T. M., Tank, D. W. & Wang, S. S. H. Widespread state-dependent shifts in cerebellar activity in locomoting mice. PLoS One 7, (2012).

9. Rancz, E. a et al. High-fidelity transmission of sensory information by single cerebellar mossy fibre boutons. Nature 450, 1245–8 (2007).

10. Chadderton, P., Margrie, T. W. & Häusser, M. Integration of quanta in cerebellar granule cells during sensory processing. Nature 428, 856–60 (2004).

11. Jörntell, H. & Ekerot, C.-F. Properties of somatosensory synaptic integration in cerebellar granule cells in vivo. J. Neurosci. 26, 11786–97 (2006).

12. Powell, K., Mathy, A., Duguid, I. & Häusser, M. Synaptic representation of locomotion in single cerebellar granule cells. Elife 4, 1–18 (2015).

13. Chen, S., Augustine, G. J. & Chadderton, P. Serial processing of kinematic signals by cerebellar circuitry during voluntary whisking. Nat. Commun. 8, 1–13 (2017).

14. Doussau, F. et al. Frequency-dependent mobilization of heterogeneous pools of synaptic vesicles shapes presynaptic plasticity. Elife 6, 1–24 (2017).

15. Valera, A. M., Doussau, F., Poulain, B., Barbour, B. & Isope, P. Adaptation of granule cell to Purkinje cell synapses to high-frequency transmission. J. Neurosci. 32, 3267–80 (2012).

16. Atluri, P. P. & Regehr, W. G. Determinants of the time course of facilitation at the granule cell to Purkinje cell synapse. J. Neurosci. 16, 5661–71 (1996).

17. Dittman, J. S., Kreitzer, a C. & Regehr, W. G. Interplay between facilitation, depression, and residual calcium at three presynaptic terminals. J. Neurosci. 20, 1374–85 (2000).

18. Miki, T. et al. Actin- and Myosin-Dependent Vesicle Loading of Presynaptic Docking Sites Prior to Exocytosis. Neuron 91, 808–823 (2016).

19. Bao, J., Reim, K. & Sakaba, T. Target-dependent feedforward inhibition mediated by short-term synaptic plasticity in the cerebellum. J. Neurosci. 30, 8171–9 (2010).

20. Zheng, N. & Raman, I. M. Synaptic inhibition, excitation, and plasticity in neurons of the cerebellar nuclei. Cerebellum 9, 56–66 (2010).

21. Brachtendorf, S., Eilers, J. & Schmidt, H. A use-dependent increase in release sites drives facilitation at calretinin-deficient cerebellar parallel-fiber synapses. Front. Cell. Neurosci. 9, 27 (2015).

22. Anwar, H., Li, X., Bucher, D. & Nadim, F. Functional roles of short-term synaptic plasticity with an emphasis on inhibition. Current Opinion in Neurobiology 43, 71–78 (2017).

23. Fioravante, D. & Regehr, W. G. Short-term forms of presynaptic plasticity. Curr. Opin. Neurobiol. 21, 269–74 (2011).

24. Grangeray-Vilmint, A., Valera, A. M., Kumar, A. & Isope, P. Short term plasticity combines with excitation-inhibition balance to expand cerebellar Purkinje cell dynamic range. J. Neurosci. (2018). doi:10.1523/JNEURØSCI.3270-17.2018

25. Sultan, F. & Bower, J. M. Quantitative Golgi study of the rat cerebellar molecular layer interneurons using principal component analysis. J. Comp. Neurol. 393, 353–373 (1998).

26. Palay, S. L. & Chan-Palay, V. Cerebellar Cortex. (Springer Berlin Heidelberg, 1974). doi:10.1007/978-3-642-65581-4

27. Sotelo, C. Molecular Layer Interneurons of the Cerebellum: Developmental and Morphological Aspects. Cerebellum 14, 534–556 (2015).

28. Rakic, P. Extrinsic cytological determinants of basket and stellate cell dendritic pattern in the cerebellar molecular layer. J. Comp. Neurol. 146, 335–354 (1972).

29. Soler-Llavina, G. J. & Sabatini, B. L. Synapse-specific plasticity and compartmentalized signaling in cerebellar stellate cells. Nat. Neurosci. 9, 798–806 (2006).

30. Bender, V. a, Pugh, J. R. & Jahr, C. E. Presynaptically expressed long-term potentiation increases multivesicular release at parallel fiber synapses. J. Neurosci. 29, 10974–8 (2009).

31. Cesca, F., Baldelli, P., Valtorta, F. & Benfenati, F. The synapsins: key actors of synapse function and plasticity. Prog. Neurobiol. 91, 313–48 (2010).

32. Song, S. & Augustine, G. J. Synapsin Isoforms and Synaptic Vesicle Trafficking. Mol. Cells 38, 936–940 (2015).

33. Malagon, G., Miki, T., Llano, I., Neher, E. & Marty, A. Counting Vesicular Release Events Reveals Binomial Release Statistics at Single Glutamatergic Synapses. Journal of Neuroscience 36, 4010–4025 (2016).

34. Nahir, B. & Jahr, C. E. Activation of extrasynaptic NMDARs at individual parallel fiber-molecular layer interneuron synapses in cerebellum. J. Neurosci. 33, 16323–33 (2013).

35. Humeau, Y., Candiani, S., Ghirardi, M., Poulain, B. & Montarolo, P. Functional roles of synapsin: lessons from invertebrates. Semin. Cell Dev. Biol. 22, 425–33 (2011).

36. Bragina, L., Giovedì, S., Barbaresi, P., Benfenati, F. & Conti, F. Heterogeneity of glutamatergic and GABAergic release machinery in cerebral cortex: analysis of synaptogyrin, vesicle-associated membrane protein, and syntaxin. Neuroscience 165, 934–943 (2007).

37. Patton, a. P., Chesham, J. E. & Hastings, M. H. Combined Pharmacological and Genetic Manipulations Unlock Unprecedented Temporal Elasticity and Reveal Phase-Specific Modulation of the Molecular Circadian Clock of the Mouse Suprachiasmatic Nucleus. J. Neurosci. 36, 9326–9341 (2016).

38. Wei, H., Masterson, S. P., Petry, H. M. & Bickford, M. E. Diffuse and specific tectopulvinar terminals in the tree shrew: Synapses, synapsins, and synaptic potentials. PLoS One 6, (2011).

39. Song, S.-H. & Augustine, G. J. Synapsin Isoforms Regulating GABA Release from Hippocampal Interneurons. J. Neurosci. 36, 6742–57 (2016).

40. Gitler, D. et al. Different presynaptic roles of synapsins at excitatory and inhibitory synapses. J. Neurosci. 24, 11368–80 (2004).

41. Feliciano, P., Matos, H., Andrade, R. & Bykhovskaia, M. Synapsin II Regulation of GABAergic Synaptic Transmission is Dependent on Interneuron Subtype. J. Neurosci. 37, 0844–16 (2017).

42. Kielland, A., Erisir, A., Walaas, S. I. & Heggelund, P. Synapsin utilization differs among functional classes of synapses on thalamocortical cells. J. Neurosci. 26, 5786–93 (2006).

43. Fassio, A., Raimondi, A., Lignani, G., Benfenati, F. & Baldelli, P. Synapsins: from synapse to network hyperexcitability and epilepsy. Semin. Cell Dev. Biol. 22, 408–15 (2011).

44. Ketzef, M. & Gitler, D. Epileptic synapsin triple knockout mice exhibit progressive longterm aberrant plasticity in the entorhinal cortex. Cereb. Cortex 24, 996–1008 (2014).

45. Hioki, H. et al. Differential distribution of vesicular glutamate transporters in the rat cerebellar cortex. Neuroscience 117, 1–6 (2003).

46. Zander, J.-F. et al. Synaptic and Vesicular Coexistence of VGLUT and VGAT in Selected Excitatory and Inhibitory Synapses. J. Neurosci. 30, 7634–7645 (2010).

47. Carter, a G. & Regehr, W. G. Prolonged synaptic currents and glutamate spillover at the parallel fiber to stellate cell synapse. J. Neurosci. 20, 4423–4434 (2000).

48. Barbour, B. Synaptic currents evoked in purkinje cells by stimulating individual granule cells. Neuron 11, 759–769 (1993).

49. Carter, A. G. & Regehr, W. G. Quantal events shape cerebellar interneuron firing. Nat. Neurosci. 5, 1309–1318 (2002).

50. Markram, H., Wang, Y. & Tsodyks, M. Differential signaling via the same axon of neocortical pyramidal neurons. Proc. Natl. Acad. Sci. 95, 5323–8 (1998).

51. Blackman, A. V., Abrahamsson, T., Costa, R. P., Lalanne, T. & Sjöström, P. J. Targetcell-specific short-term plasticity in local circuits. Frontiers in Synaptic Neuroscience 5, 1–13 (2013).

52. Koester, H. J. & Johnston, D. Target cell-dependent normalization of transmitter release at neocortical synapses. Science 308, 863–6 (2005).

53. Scanziani, M., Gähwiller, B. H. & Charpak, S. Target cell-specific modulation of transmitter release at terminals from a single axon. Proc. Natl. Acad. Sci. 95, 12004–9 (1998).

54. Delaney, A. J. & Jahr, C. E. Kainate receptors differentially regulate release at two parallel fiber synapses. Neuron 36, 475–82 (2002).

55. Pelkey, K. a, Topolnik, L., Lacaille, J.-C. & McBain, C. J. Compartmentalized Ca(2+) channel regulation at divergent mossy-fiber release sites underlies target cell-dependent plasticity. Neuron 52, 497–510 (2006).

56. Buchanan, K. a. et al. Target-Specific Expression of Presynaptic NMDA Receptors in Neocortical Microcircuits. Neuron 75, 451–466 (2012).

57. Galvan, E. J. & Gutierrez, R. Target Dependent Compartmentalization of the Corelease of Glutamate and GABA from the Mossy Fibers. J. Neurosci. 37, 701–714 (2016).

58. Espinosa, J. S. & Luo, L. Timing neurogenesis and differentiation: insights from quantitative clonal analyses of cerebellar granule cells. J. Neurosci. 28, 2301–12 (2008).

59. Wilms, C. D. & Häusser, M. Reading out a spatiotemporal population code by imaging neighbouring parallel fibre axons in vivo. Nat. Commun. 6, 6464 (2015).

60. Owald, D. & Sigrist, S. J. Assembling the presynaptic active zone. Current Opinion in Neurobiology 19, 311–318 (2009).

61. Brenowitz, S. D. & Regehr, W. G. Reliability and heterogeneity of calcium signaling at single presynaptic boutons of cerebellar granule cells. J. Neurosci. 27, 7888–98 (2007).

62. Zhang, W. & Linden, D. J. Calcium influx measured at single presynaptic boutons of cerebellar granule cell ascending axons and parallel fibers. Cerebellum 11, 121–131 (2012).

63. Zhang, W. & Linden, D. J. Neuromodulation at single presynaptic boutons of cerebellar parallel fibers is determined by bouton size and basal action potential-evoked Ca transient amplitude. J. Neurosci. 29, 15586–94 (2009).

64. Bouvier, G. et al. Burst-Dependent Bidirectional Plasticity in the Cerebellum Is Driven by Presynaptic NMDA Receptors. Cell Rep. 104–116 (2016). doi:10.1016/j.celrep.2016.03.004

65. Beierlein, M. & Regehr, W. G. Local interneurons regulate synaptic strength by retrograde release of endocannabinoids. J. Neurosci. 26, 9935–43 (2006).

66. Schmidt, H. et al. Nanodomain coupling at an excitatory cortical synapse. Curr. Biol. 23, 244–9 (2013).

67. Kusch, V. et al. Munc13-3 Is Required for the Developmental Localization of Ca 2+ Channels to Active Zones and the Nanopositioning of Ca v 2.1 Near Release Sensors. Cell Rep. 22, 1965–1973 (2018).

68. Ishiyama, S., Schmidt, H., Cooper, B. H., Brose, N. & Eilers, J. Munc13-3 Superprimes Synaptic Vesicles at Granule Cell-to-Basket Cell Synapses in the Mouse Cerebellum. J. Neurosci. 34, 14687–96 (2014).

69. Südhof, T. C. Neurotransmitter release: the last millisecond in the life of a synaptic vesicle. Neuron 80, 675–90 (2013).

70. Giovedì, S. et al. Synapsin is a novel Rab3 effector protein on small synaptic vesicles. I. Identification and characterization of the synapsin I-Rab3 interactions in vitro and in intact nerve terminals. J. Biol. Chem. 279, 43760–8 (2004).

71. Medrihan, L. et al. Synapsin II desynchronizes neurotransmitter release at inhibitory synapses by interacting with presynaptic calcium channels. Nat. Commun. 4, 1512 (2013).

72. Hirano, M. et al. C-terminal splice variants of P/Q-type Ca 2+ channel Ca V 2.1 α 1 subunits are differentially regulated by Rab3-interacting molecule proteins. J. Biol. Chem. 292, 9365–9381 (2017).

73. Kintscher, M., Wozny, C., Johenning, F. W., Schmitz, D. & Breustedt, J. Role of RIM1α in short- and long-term synaptic plasticity at cerebellar parallel fibres. Nat. Commun. 4, 2392 (2013).

74. Chadderton, P., Schaefer, A. T., Williams, S. R. & Margrie, T. W. Sensory-evoked synaptic integration in cerebellar and cerebral cortical neurons. Nat. Rev. Neurosci. 15, 71–83 (2014).

75. Arenz, A., Silver, R. A., Schaefer, A. T. & Margrie, T. W. The contribution of single synapses to sensory representation in vivo. Science 321, 977–80 (2008).

76. Chabrol, F. P., Arenz, A., Wiechert, M. T., Margrie, T. W. & DiGregorio, D. a. Synaptic diversity enables temporal coding of coincident multisensory inputs in single neurons. Nat. Neurosci. 18, (2015).

77. Kistler, W. M., van Hemmen, J. L. & De Zeeuw, C. I. Time window control: a model for cerebellar function based on synchronization, reverberation, and time slicing. Prog. Brain Res. 124, 275–97 (2000).

78. Mauk, M. D. & Buonomano, D. V. THE NEURAL BASIS OF TEMPORAL PROCESSING. Annu. Rev. Neurosci. 27, 307–340 (2004).

79. Wolpert, D. M., Miall, R. C. & Kawato, M. Internal models in the cerebellum. Trends in Cognitive Sciences 2, 338–347 (1998).

80. Binda, F. et al. Inhibition promotes long-term potentiation at cerebellar excitatory synapses. Sci. Rep. 6, 33561 (2016).

81. Suvrathan, A., Payne, H. L. & Raymond, J. L. Timing Rules for Synaptic Plasticity Matched to Behavioral Function. Neuron 92, 959–967 (2016).

82. Rosahl, T. W. et al. Essential functions of synapsins I and II in synaptic vesicle regulation. Nature 375, 488–93 (1995).

83. Valera, A. M. et al. Stereotyped spatial patterns of functional synaptic connectivity in the cerebellar cortex. Elife 5, 1–22 (2016).

84. Longair, M. H., Baker, D. a. & Armstrong, J. D. Simple neurite tracer: Open source software for reconstruction, visualization and analysis of neuronal processes. Bioinformatics 27, 2453–2454 (2011).

85. Korogod, N., Petersen, C. C. H. & Knott, G. W. Ultrastructural analysis of adult mouse neocortex comparing aldehyde perfusion with cryo fixation. Elife 4, 1–17 (2015).

86. Palay, S. L. & Chan-Palay, V. A guide to the synaptic analysis of the neuropil. Cold Spring Harbor Symposia on Quantitative Biology 40, 1–16 (1976).

